# FAM57B is a modulator of ceramide synthesis that regulates sphingolipid homeostasis and synaptic composition in the developing brain

**DOI:** 10.1101/2021.03.02.433619

**Authors:** Danielle L. Tomasello, Jiyoon L. Kim, Yara Khodour, Jasmine M. McCammon, Maya Mitalipova, Rudolf Jaenisch, Anthony H. Futerman, Hazel Sive

**Author notes:** To whom correspondence should be addressed at: 115 Richards Hall, 360 Huntington Ave, Boston MA 02115, USA. Northeastern University College of Science, Boston, MA 02115.

## Abstract

The complex 16p11.2 Deletion Syndrome (16pdel) is accompanied by neurological disorders, including epilepsy, autism spectrum disorder and intellectual disability. We demonstrate that 16pdel iPSC differentiated neurons showed augmented local field potential activity and altered ceramide-related lipid species relative to unaffected. *FAM57B*, a poorly characterized gene in the 16p11.2 interval, has emerged as a candidate tied to symptomatology. We found that FAM57B modulates ceramide synthase (CerS) activity, but is not a CerS per se. In *FAM57B* mutant human neuronal cells and zebrafish brain, composition and levels of sphingolipids and glycerolipids associated with cellular membranes are disrupted. Consistently, we observed aberrant plasma membrane architecture and synaptic protein mislocalization, which were accompanied by depressed brain and behavioral activity. Together, these results suggest that haploinsufficiency of *FAM57B* contributes to changes in neuronal activity and function in 16pdel Syndrome, through a crucial role for the gene in lipid metabolism.

## Main

16p11.2 Deletion (16pdel) Syndrome, a severe and prevalent neurodevelopmental disorder, is a copy number variant with deletion of ∼600 kb from chromosome 16. This haploinsufficiency syndrome is estimated to affect ∼1 in 2500 worldwide and is tightly associated with autism spectrum disorder (ASD), language and intellectual disability, seizures, attention-deficit/hyperactivity disorder, macrocephaly, hypotonia and obesity^1–5^. Strong indications of synaptic defects are associated with 16pdel symptoms, particularly epilepsy^6,7^ and ASD^6,8- 13^, as well as links to metabolic defects^14^.

Previously, analysis in the zebrafish model suggested that *FAM57B* is a pivotal hub gene in the 16p11.2 interval, that encodes a protein proposed to be a ceramide synthase^15^. Using a pairwise partial loss of function screen for zebrafish embryonic brain morphology, we found that *fam57b* interacted with numerous other 16p11.2 interval genes, suggesting haploinsufficiency of *FAM57B* is critical in 16pdel Syndrome etiology^16^. FAM57B (family with sequence similarity 57, member B) is a Tram-Lag-CLN8 (TLC) family member, containing a domain of roughly 200 amino acids found in ceramide synthases (CerS)^17^. Ceramides are sphingolipids (SLs) which are key membrane components and also act as signaling molecules to modulate proliferation, apoptosis, inflammation, cell cycle arrest and ER stress^18^. In humans, mutations in some of the 6 known CerS are associated with autism, epilepsy and intellectual disability^19–21^. In this study, to further assess the predicted connection with 16pdel Syndrome, we examined FAM57B function through a multidisciplinary approach, across human cells and the zebrafish system.

## Results

### Augmented network activity in 16pdel neuron cultures

Based on previous data, we hypothesized that 16pdel neurons would show an altered lipid profile due to contributions of *FAM57B* and possibly other 16pdel genes with predicted roles in metabolism^16^. To test this, we prepared neurons from 16pdel carrier induced-pluripotent stem cells (iPSC), part of the Simons VIP Consortium, and unaffected control iPSC in culture^22^ (**Supp Table 1**). Neural progenitor cells were differentiated into cortical neurons, since the cortex has consistently shown anatomical differences in 16pdel affected individuals^9,23–27^. After one month in culture, immunocytochemistry (ICC) indicated mature neurons by presence of vesicular glutamate 1 and 2 receptors (VGlut1/2) (**Supp Figs. 1a,b**), the synaptic markers PSD95 (**Supp Fig. 1b**) and Synaptotagmin-1 (Syt1). Cultures of control and 16pdel (proband) differentiated neurons showed similar percentages of mature neurons by these criteria.

**Figure 1.**
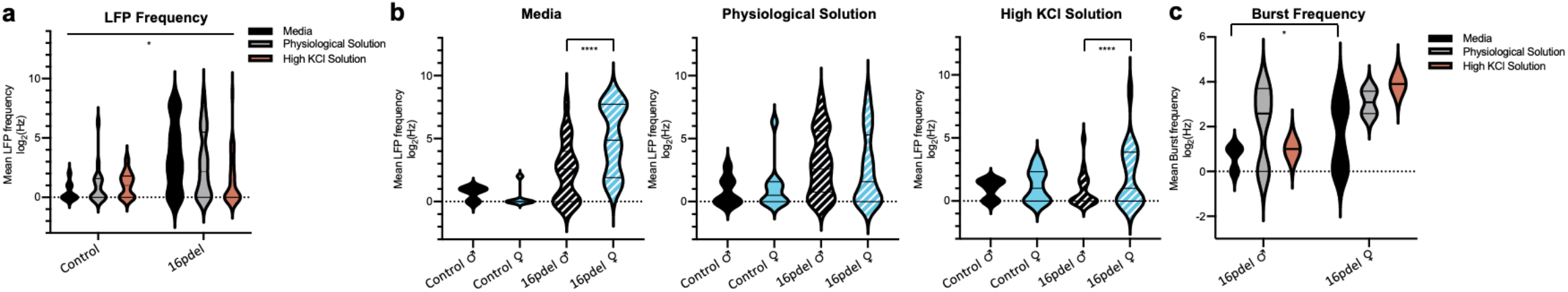
Augmented local field potential activity in 16pdel syndrome differentiated neuronal culture. **a)** Local Field Potential (LFP) summary analyzed by log2 fold change between control and 16pdel patient differentiated neurons. MEA activity was recorded over 30 min starting in media, followed by physiological and high potassium chloride (KCl) solution. Data was summarized and pooled from 3 experiments. Control n = 9 (media), n = 17 (Physiological Solution), n = 13 (High KCl Solution). 16pdel n = 35 (media), n = 76 (Physiological Solution), n = 35 (High KCl Solution). Violin plot group analysis: Control −16pdel 2-way ANOVA. *p ≤ 0.05 **b)** Increased sex specific activity in female 16pdel probands drives overall increased LFPs, compared to unaffected controls. Media unaffected neurons (Control) male (♂) n = 3, Control female (♀) n = 6,16pdel neurons (Proband) ♂ n = 21, Proband ♀ n = 14, Physiological Solution Control ♂ n = 7, Control ♀ n = 10, Proband ♂ n = 38, Proband ♀ n = 38, High KCl Solution Control ♂ n = 6, Control ♀ n = 7, Proband ♂ n = 18, Proband ♀ n = 17. Violin plot analysis: male vs female T-Test. ****p ≤ 0.0001. **c)** Increased sex specific female electrogenic burst frequency analyzed by log2 fold change between 16pdel male and female Media ♂ n = 21, ♀ n = 14, Physiological Solution ♂ n = 38, ♀ n = 38, High KCl Solution ♂ = 18, ♀ n = 17. Violin plot analysis: male vs female T-Test. *p ≤ 0.05.

To further characterize these neurons, we probed network electrical activity by multi-electrode array (MEA). Spontaneous activity of differentiated neurons was measured over thirty minutes; first in culture media, followed by physiological solution and last in high potassium chloride solution. Comparing grouped genotypes, we recorded an increased frequency of Local Field Potentials (LFPs) in 16pdel proband neuron cultures relative to controls, indicating 16pdel neurons display heightened spontaneous and evoked activity compared to unaffected control (**Fig. 1a**). Examining individual patient cell lines, we observed relatively similar MEA activity in controls (black), and increased electrical activity in 16pdel neuron cultures (grey) (**Supp Fig. 2a**). Interestingly, female 16pdel neuron cultures showed statistically increased LFP frequency compared to male 16pdel neuron cultures when measured in media and High KCl solution (**Fig. 1b**). Sex differences were also observed in LFP firing and bursting properties, with increased burst frequency of female 16pdel neurons compared to male in media (**Fig. 1c**). Our analysis shows differences between activity of 16pdel and control neurons. These findings expand previous observations that demonstrated larger cell size and deficits in synaptic density in 16pdel neurons compared to control^28^.

**Figure 2.**
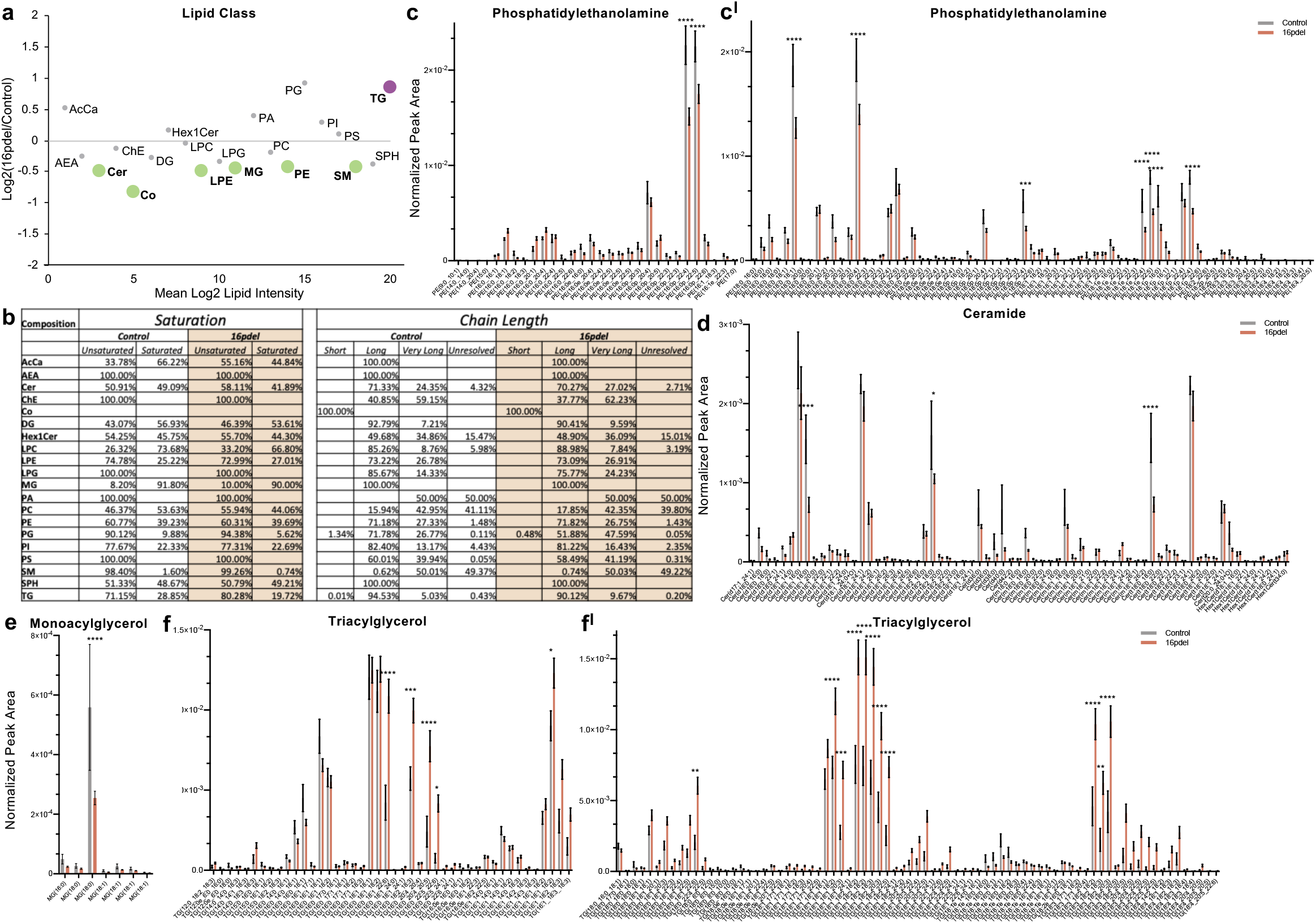
Significant lipid changes between control and 16pdel differentiated neurons. **a)** Total log2 fold change from normalized peak area of lipid class analysis from untargeted lipidomics. Bolded and colored indicate statistically significant changes by T-Test, p ≤ 0.05 −0.0001. AcCa acyl carnitine, AEA N-arachidonoylethanolamine, Cer ceramide, ChE cholesterol ester, Co coenzyme, DG diacylglycerol, Hex1Cer hexosylceramide, LPC lysophosphatiylcholine, LPE lysophosphatiylethanolamine, LPG lysophosphatiylglycerol, MG monoacylglycerol, PA phosphatidic acid, PC phosphatidylcholine, PE phosphatidylethanolamine, PG phosphatidylglycerol, PI phosphatidylinositol, PS phosphatidylserine, SM sphingomyelin, SPH sphingosine,TG triacylglycerol. **b)** Total lipid composition analysis from untargeted lipidomics between control and 16pdel neuron. Chain Length: Small 1-5, Medium 6-12, Long 13-21, Very Long 22+, and Unresolved. **c – f)** Selected analysis of lipid species from untargeted lipidomics classes. Lipid Class specified for each histogram (c -phosphatidylethanolamine, d -ceramide, e -monoacylglycerol, f – triacylglycerol) normalized peak area between control (grey) and 16pdel (orange). Statistical analysis by 2-Way ANOVA, *p ≤ 0.05 **p ≤ 0.01,***p ≤ 0.001, ****p ≤ 0.0001. Control n = 10, 16pdel n= 69, error bars represent SEM. TG and PE long and very long chain species not shown as no significant differences were found by ANOVA.

### Significant lipid class and individual species changes indicates complexity of 16pdel Syndrome

Using differentiated 16pdel and control neurons, we compared their lipid cohorts using untargeted lipidomics. Consistent with predictions, many significant changes were observed in total lipid classes between 16pdel and control neurons (**Fig. 2a**). Levels of SLs (ceramide (Cer) and sphingomyelin (SM)) and glycerolipids (GL) (lysophosphatidylethanolamine (LPE), phosphatidylethanolamine (PE), monoacylglycerol (MG)) were significantly decreased, while GL (triacylglycerol (TG)) levels significantly increased. The inverse changes between MG and TG suggests addition or removal of one acyl chain. Analyzing lipid composition, we found similar levels of unsaturated and saturated species between 16pdel and control neurons, but differences in saturation of acyl carnitine (AcCa) (unsaturated control 33.78% vs 16pdel 55.16%, and saturated control 66.22% and 16pdel 44.84%) and TG (unsaturated control 71.15% vs 16pdel 80.28%, and saturated control 28.85% and 16pdel 19.72%) (**Fig. 2b**). Polyunsaturated fatty acids (PUFAs) are important in the brain, where they are essential for signaling and membrane structure^29^. Chain length analysis indicated large differences in lysophosphatidylglycerol (LPG) (long chain control 85.76% vs 16pdel 75.55%) and phosphatidylglycerol (PG) (long chain control 71.78% vs 16pdel 51.88%). While having a similar ratio of long and very long chain PE species (**Fig. 2b**), analysis of individual lipid species demonstrated significantly decreased levels of several PE(18:22) species in 16pdel neurons relative to control (**Fig. 2c**). Additionally, decreased Cer(18) species were observed in 16pdel (**Fig. 2d**). Comparing MG and TG, MG(18:0) decreased while many TG(18:1,18:2,18:3) increased (**Fig. 2e,f**). Together, this analysis identifies differences in metabolism of ceramides and GLs in 16pdel neurons that are critical for function of the ER, mitochondria and plasma membrane^30^. The shift in saturation and tail length of GLs between 16pdel and control neurons suggests dysfunctional neuronal membrane.

### FAM57B functions as a ceramide synthase modulator

We considered that the extensive lipid differences between 16pdel and control neurons may partly result from FAM57B activity. The function of this protein is not clear, although a single report suggests that FAM57B has ceramide synthase activity^15^. However, our sequence analysis indicates that while FAM57B is part of the TLC protein family, including ceramide synthases (CerS), FAM57B lacks seven highly conserved residues (DxRSDxE) that may distinguish CerS from other members of the TLC familly^17^ (**Supp Fig. 3)**. To determine whether FAM57B is a *bone fide* CerS, it was expressed in *CerS2*^*-/-*^ *(KO)* HEK293T cells, which lack endogenous CerS2 activity^31^ (**Fig. 3a**). No CerS2 activity was detected in *CerS2 KO* cells upon transfection of FAM57B alone. However, co-transfection of FAM57B with CerS2 resulted in a significant increase in CerS2 activity compared to transfection of CerS2 alone (**Fig. 3a**), suggesting that FAM57B might modulate CerS2 activity. There are six CerS isoforms in mammals, where each uses a restricted subset of acyl CoAs of defined chain length for ceramide synthesis^32^. To assess whether FAM57B can modulate other members of the mammalian CerS family; we expressed CerS5 and CerS6 with or without co-transfection of FAM57B in wildtype HEK293T cells. Upon co-transfection of CerS2 with FAM57B in wildtype HEK293T cells, levels of CerS2 activity and expression were significantly increased compared to CerS2 alone (**Fig. 3b**). While co-transfection of FAM57B with CerS5 did not alter expression nor activity of this CerS **(Fig. 3c)**, an opposite trend was seen upon co-transfection of FAM57B with CerS6, whose activity decreased upon co-transfection with FAM57B (**Fig. 3c**). These results suggest that FAM57B affect protein levels and activity of certain CerS isoforms, and may do so by an indirect mechanism, dependent on interaction of the two proteins. This hypothesis was confirmed by immunoprecipitation, in which Flag-tagged FAM57B was able to interact with all three HA-tagged CerS isoforms (**Fig. 3d**). These data newly implicate FAM57B as a modulator of CerS, but refute a previous report that this protein functions as a CerS^15^.

**Figure 3.**
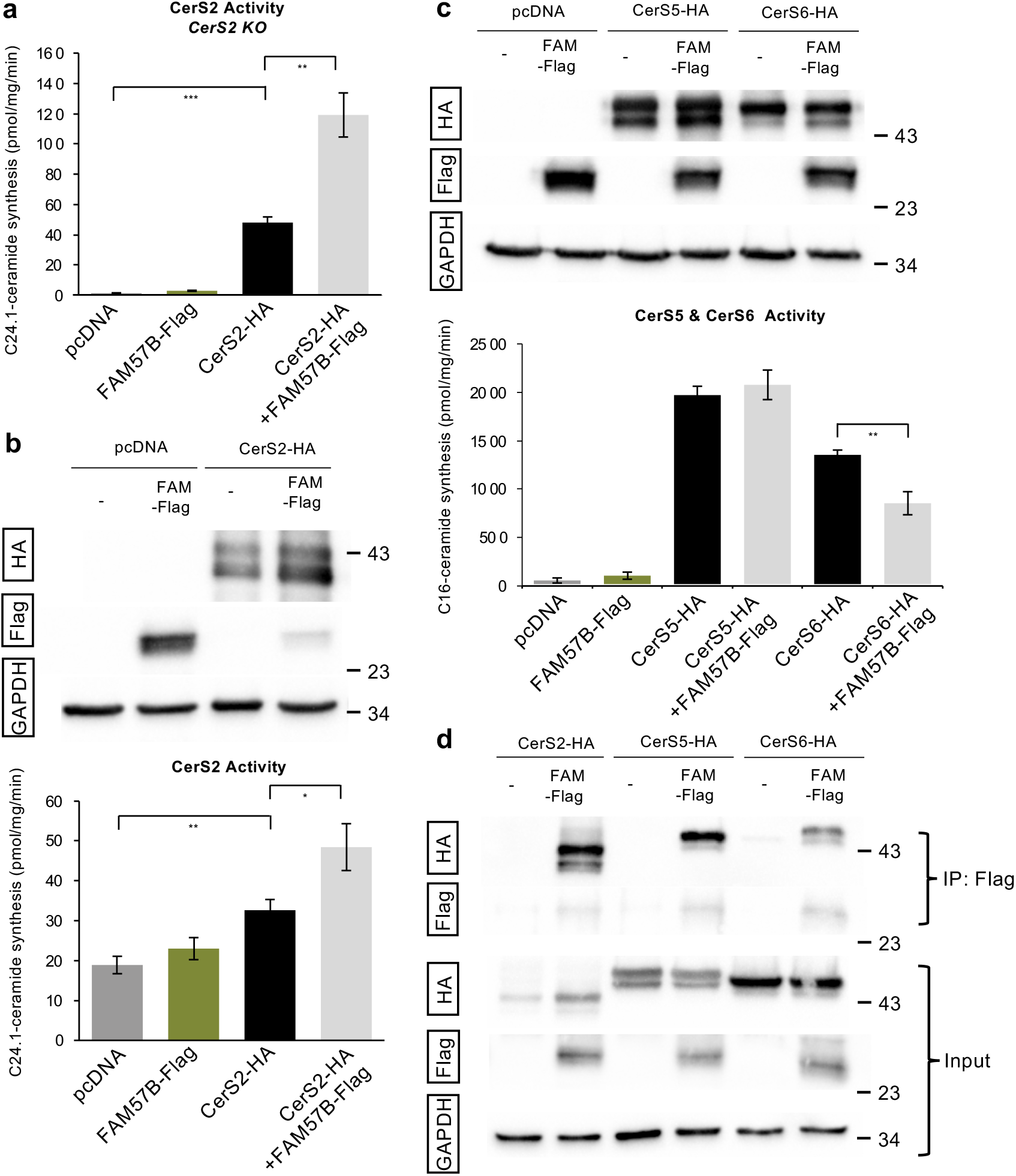
FAM57B interacts with CerS but does not have CerS activity. **a)** CerS2 activity assayed using C24:1-CoA in *CerS2 KO* HEK293T cells. Statistical analysis by T-test *p ≤ 0.05, **p ≤ 0.01, error bars SEM. **b)** (Upper) Western blot analysis of total human FAM57B-Flag and CerS2-HA after transfection in HEK293T cells. Proteins were prepared from HEK293T cells overexpressing the indicated constructs. Anti-HA and anti-Flag are indicated. (Lower) CerS2 activity assayed using C24:1-CoA in HEK293T cells. GAPDH was used as a loading control. Statistical analysis by T-test *p ≤ 0.05, **p ≤ 0.01, error bars SEM. **c)** (Upper) Western blot analysis of total human FAM57B-Flag, CerS5-HA and CerS6-HA after transfection in HEK293T cells. Proteins were prepared from cells overexpressing the indicated constructs. Anti-HA and anti-Flag are indicated. (Lower) CerS5 and CerS6 activity was assayed using C16:0-CoA in HEK293T cells. Anti-HA and anti-Flag are indicated. GAPDH was used as a loading control. **d)** Total cell lysates were prepared from the co-transfected cells with FAM57B-Flag and CerS2, 5 or 6-HA constructs and solubilized with 1% NP-40. Total lysates (input) or proteins immuno-precipitated with anti-Flag M2 agarose (IP) were subjected to immunoblotting with anti-HA or anti-Flag antibodies. GAPDH was used as a loading control.

### *FAM57B* modulates lipid cohorts and synaptic proteins in human cells

The intriguing functional differences between 16pdel and control neurons raises the question of whether *FAM57B* haploinsufficiency contributes to these differences. To address this, we used the human neuroblastoma line SH-SY5Y to create a knockout (*FAM57B KO*) and *FAM57B* heterozygote (*FAM57B HET*) using CRISPR-Cas9 editing. SH-SY5Y cells have proven useful for studying neuronal properties and function^33^. For our studies, SH-SY5Y cells were differentiated into neurons after incubation in media containing retinoic acid. Overall, total lipid classes showed significant differences between *FAM57B KO* and WT (wildtype), specifically, increased ChE and MG (**Fig. 4a**). Comparing *FAM57B HET* to WT, we observed increased LPC (**Fig. 4b**). Additionally, relative to *FAM57B HET*, we found HexCer and PG significantly decreased while ChE increased in *FAM57B KO* cells (**Fig. 4c**).

**Figure 4.**
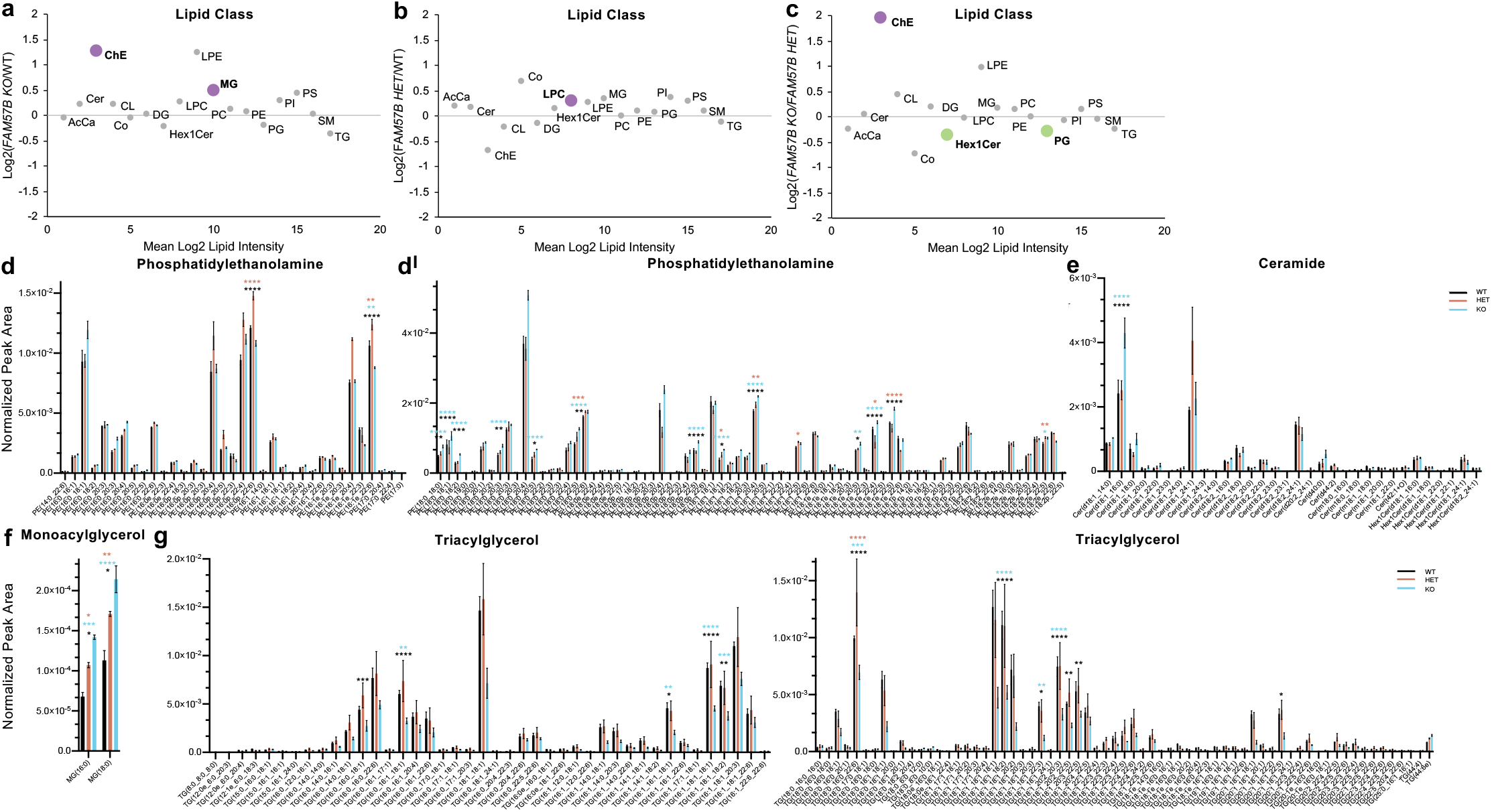
Significant lipid changes in sphingolipids and glycerolipids between WT and *FAM57B mutant* human differentiated SH-SY5Y neuronal cells. **a -c)** Total log2 fold change from normalized peak area of lipid class analysis from untargeted lipidomics. **a)** *FAM57B KO* – WT, **b)** *FAM57B HET* – WT, **c)** *FAM57B KO* – *FAM57B HET*. Bolded and colored indicate statistically significant changes by T-Test, p ≤ 0.05 −0.0001. AcCa acyl carnitine, Cer ceramide, ChE cholesterol ester, CL cardiolipin, Co coenzyme, DG diacylglycerol, HexCer Hexosylceramide, LPC lysophosphatiylcholine, LPE lysophosphatiylethanolamine, MG monoacylglycerol, PC phosphatidylcholine, PE phosphatidylethanolamine, PG phosphatidylglycerol, PI phosphatidylinositol, PS phosphatidylserine, SM sphingomyelin, TG triacylglycerol. **d – g)** Selected analysis of lipid species from untargeted lipidomics classes. Lipid Class specified for each histogram, normalized peak area between WT (black) *FAM57B HET* (orange) and *FAM57B KO* (blue). Statistical analysis by 2-Way ANOVA, *p ≤ 0.05 **p ≤ 0.01, ***p ≤ 0.001, ****p ≤ 0.0001. Color of asterisks indicate comparison between WT – HET (orange), WT – KO (blue), HET – KO (black). WT n = 3, *FAM57B HET* n= 3, *FAM57B KO* n = 3, error bars represent SEM. Experiment repeated twice, analysis was similar between two separate runs.

Notably, lipid class differences between *FAM57B KO, FAM57B HET* and WT were also significantly altered in 16pdel patient neurons compared to controls (**Fig. 2a**). In both *FAM57B KO* and *FAM57B HET* relative to WT, we observed increased abundance of PE(18:0,18:1,22:4,22:5) (**Fig. 4d**), Cer(d18:1) (**Fig. 4e**), MG(18:0) (**Fig. 4f**), and with decreased abundance of TG(16:0,16:1,18:0,18:1,18:2,22:6) **(Fig. 4g**). These differences are similar to those seen in 16pdel relative to control neurons **(Fig. 2c-f)**. In *FAM57B KO* relative to WT, there was increased abundance of MG and decreased abundance TG (**Fig. 4f,g**), similar to 16pdel neurons compared to control (**Fig. 2a**,**e**,**f**). The alterations in lipid cohorts between *FAM57B KO* and *FAM57B HET* human neurons is similar with lipid changes in 16pdel neurons, and consistent with a role for FAM57B in dosage-sensitive lipid regulation, including tight regulation of HexCer. The similarities between alterations in lipid cohorts between *FAM57BKO* and *FAM57B HET* human neurons and 16pdel neurons, is consistent with a role for FAM57B in dosage-sensitive, lipid regulation, including tight regulation of HexCer.

In addition to comparing lipid cohorts, we looked for other differences between neurons differentiated from WT, *FAM57B KO* and *FAM57B HET* cells. Relative to WT, *FAM57B KO* neurons showed severely decreased levels of β-Actin protein, an intermediate filament cytoskeletal protein essential for axon guidance, synaptogenesis, and extracellular structures that are connected to the plasma membrane, by Western analysis and immunocytochemistry^34^ (**Fig. 5a-b**). *FAM57B HET* showed a relative decrease of roughly half of β-Actin compared to WT, suggesting gene dosage dependent changes (**Fig. 5a-b**). β-Actin mRNA and protein localization is found to modulate cytoskeletal composition and growth cone dynamics during neuronal development^35^. These data suggest that loss of FAM57B may affect plasma membrane architecture and protein localization.

**Figure 5.**
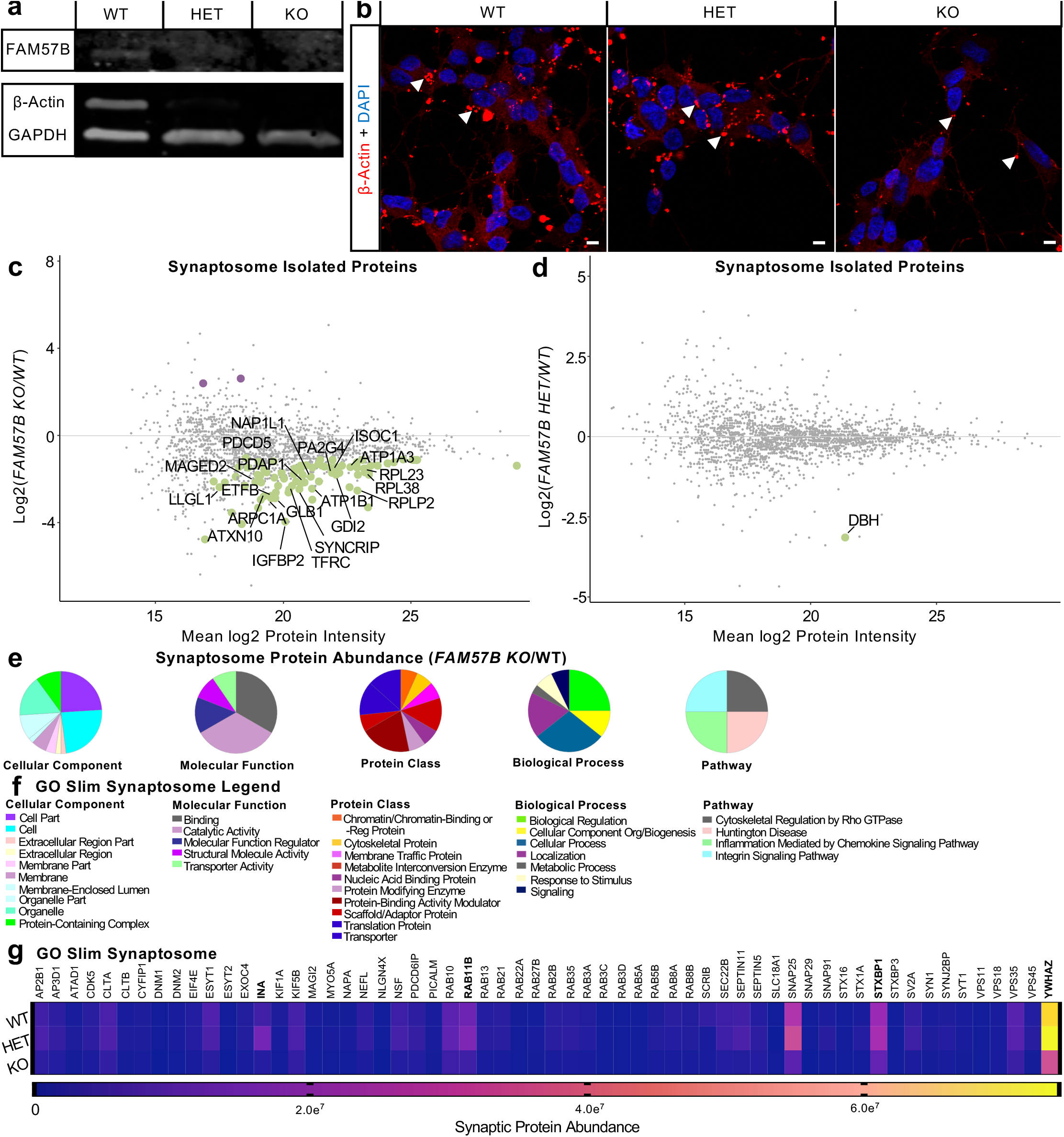
*FAM57B* knockout human neurons indicate altered cytoskeleton and diminished synaptic proteins. **a)** Validation of *FAM57B HET* and *KO* protein levels compared to WT control in SH-SY5Y RA differentiated neuronal cells. Decreased Beta-Actin expression observed in *FAM57B HET* and *KO* compared to W*T*. GAPDH used as loading control. Repeated thrice. **b)** Staining of differentiated cells with deceased Beta-Actin (arrow). Scale bar = 5 μm. Repeated twice. **c -d)** Isolated synaptosome protein abundance changes between **c)** *FAM57B KO* relative to WT and **d)** *FAM57B HET* relative to WT (Log2 Fold). Labeled and colored indicating increased (purple) or decreased (green) abundance. Only the top 20 proteins of statical significance were labeled in **c** and analyzed in **e-g**. WT n = 8, HET n = 10, KO n = 7. indicating increased (purple) or decreased (green) abundance. Lowest abundance membrane protein Synaptotagmin-1a (red box). n = 6 per genotype. **e – f)** Gene ontology analysis of statistically significant synaptosome isolated proteins (e) in *FAM57B KO* relative to WT. **e)** gene ontology pie graphs of top 20 decreased protein groups of cellular components, molecular function, protein classes, biological processes and pathways. **f)** gene ontology figure legend. **g)** Analysis of synaptic markers from isolated synaptosomes between all 3 genotypes. Significantly decreased protein abundance of synaptic structural and maturation proteins, and vesicle regulation machinery. Bolded: INA – WT -HET & WT -KO, RAB11B – HET -KO, STXBP1 – WT -HET & WT -KO, YWHAZ – WT -HET & WT -KO. 2-Way ANOVA, p ≤ 0.05 – 0.0001.

To understand the consequence of *FAM57B loss of function* on neuronal maturation and function, we probed synaptic composition. Synaptosomes, comprising the pre-and postsynaptic membranes and postsynaptic density, were isolated from unfixed cells and processed by MS/MS. Synaptosomes from *FAM57B KO* cells showed significantly decreased abundance of over 100 proteins relative to WT (**Fig. 5c**). In contrast, *FAM57B HET* showed no statistically significant changes in synaptosome protein composition relative to WT (**Fig. 5d**), with the exception of Dopamine Beta-Hydroxylase (DBH). Among the top 20 significantly decreased proteins in the *FAM57B KO* synaptosomes were those associated with protein trafficking, localization and stabilization (**Fig. 5e,f)**. Additionally, we observed an overall decrease in levels of synaptic proteins in *FAM57B KO* compared to *FAM57B HET* and *WT* (**Fig. 5g**) when analyzing hallmark proteins that function at the synapse. These decreases included α-internexin INA, small GTPase vesicle recycling RAB11B, SNARE protein syntaxin STXBP1 and scaffolding protein YWHAZ. Similar to β-Actin, INA is a neurofilament subunit protein important for neuronal cytoskeletal assembly and synaptogenesis localized to the post-synaptic terminal^36^.

Together, these data indicate that in human neurons mutant or heterozygous for *FAM57B*, there are significant changes in lipid composition, membrane dynamics and synaptic proteins. The data are consistent with the suggestion that a deficit in FAM57B function partly contributes to 16pdel neuronal anomalies relative to control. The smaller changes observed in *FAM57B HET* relative to *FAM57B KO* suggests that other 16pdel genes contribute to phenotypes in the haploinsufficient syndrome.

### FAM57B is essential for Sphingolipid (SL) and Glycerolipid (GL) homeostasis in the developing brain

16pdel alters brain structure and function, including neuroanatomical abnormalities and increased risk of psychiatric and other brain disorders^37,38^. To understand how *FAM57B* contributes to brain development, we analyzed zebrafish, *Danio rerio*, a powerful system for analysis of neural development and neurodevelopmental disorders^39–42^. The zebrafish genome includes two copies of the *fam57b* gene, *fam57ba* and *fam57bb*. We used CRISPR to build double (null) mutants, *fam57ba*^*-/-*^;*fam57bb*^*-/-*^ (*fam57b mut*), and heterozygotes *fam57b*^*+/-*^;*fam57bb*^*+/-*^ *(fam57b het)*, to assess dosage effects of FAM57B.

To determine whether *fam57b* regulates lipid metabolites, we performed untargeted lipid profiling on *fam57b mut* and *fam57b het* zebrafish brain tissue at 7 days post-fertilization (7 dpf), an optimal timepoint for molecular and behavioral studies of a developing yet complex brain^43^. Striking differences in SL and GL lipid abundances were present in *fam57b mut* and *fam57b het* compared to wildtype (AB) zebrafish brain (**Fig. 6a,b**). By lipid class, there was a significant increase in Cer, LPE, MG and SM along with phosphatidylinositol (PI) and cardiolipin (CL) and decreased PS in *fam57b mut* compared to AB (**Fig. 6a**). A similar trend to *fam57b mut*, with increased hexosylceramide (HexCer) and decreased PS lipid classes, were defined in *fam57b het* brains compared to AB control. An overlap in lipid differences were observed between *fam57b mut* brains and FAM57B KO human neurons, with increased abundance of MG class, and Cer(d18:1) and MG(18:0) species (**Figs. 6a, 4a, 6d,e, 4d,e**). Many PE species similarly increased, PE(18:0,18:1,20:4), comparing *fam57b mut* and *FAM57B KO* to controls. An important finding across all systems compared, including 16pdel syndrome patient neurons, heterozygous and mutant *FAM57B* cells and larvae brains, is a change in ether-linked PE (**Figs. 2c, 4d, 6c**,**g**). Ether GLs differ in phase-transition temperature from gel to liquid crystalline and from lamellar to hexagonal phases, and are proposed to regulate properties of neuronal membranes^44,45^. In addition to PE, ceramides were altered in *fam57b mut* and *fam57b het* brain tissue compared to AB (**Fig. 4d**,**h**). Lipidomics resolved predominantly Cer(d18:1) species in zebrafish brains, which agrees with previously published ceramide composition at 7 dpf^46^. These findings suggest a key role for Fam57b in SL and GL regulation during brain development.

**Figure 6.**
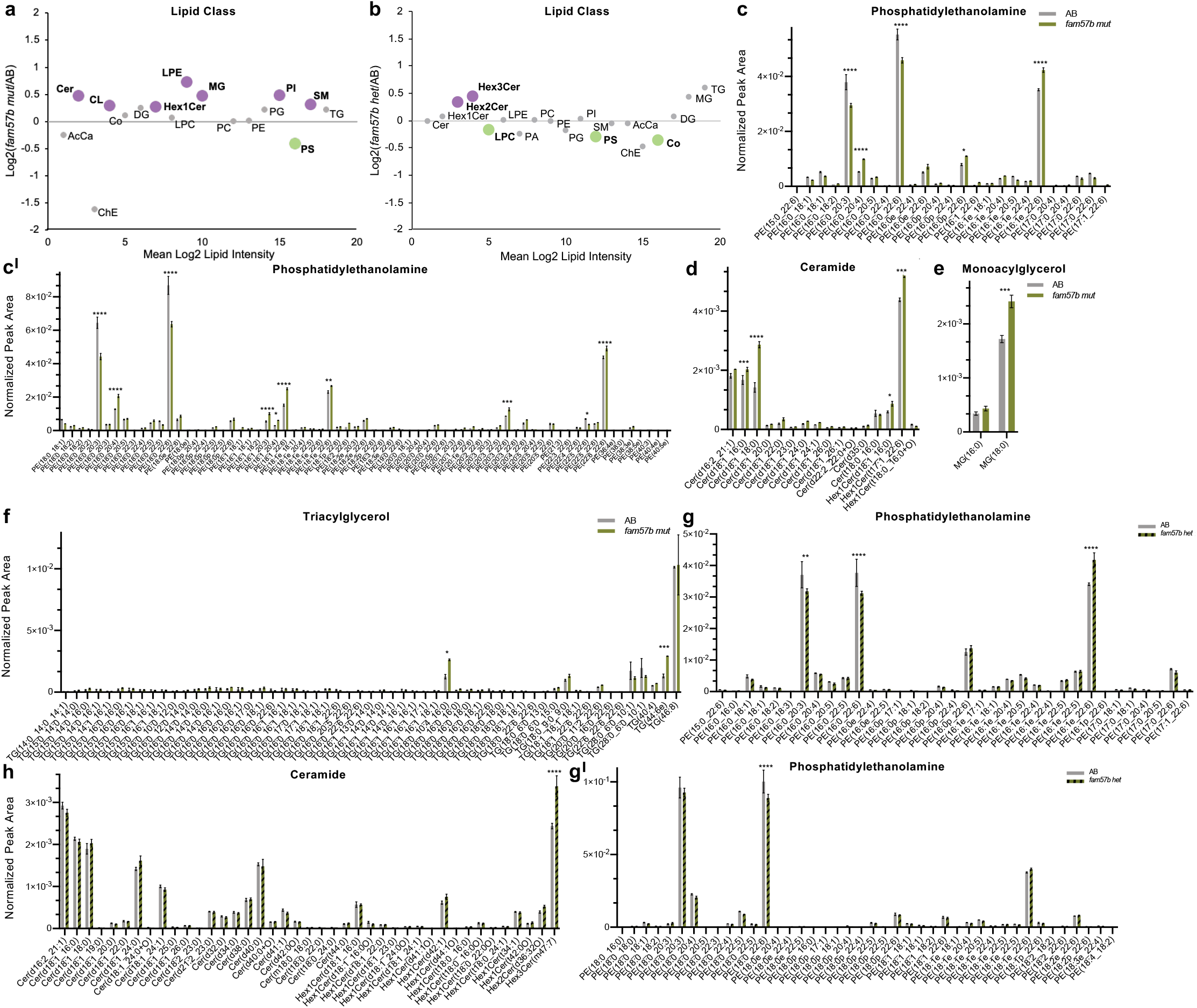
Significant lipid changes in ceramides and glycerols between AB and *fam57b mut* brain tissue. **a**) Total log2 fold change from normalized peak area of lipid class analysis from untargeted lipidomics. Bolded and colored indicate statistically significant changes by T-Test, p ≤ 0.05 −0.0001. AcCa acyl carnitine, Cer ceramide, ChE cholesterol ester, CL cardiolipin, Co coenzyme, DG diacylglycerol, HexCer Hexosylceramide, LPC lysophosphatiylcholine, LPE lysophosphatiylethanolamine, MG monoacylglycerol, PC phosphatidylcholine, PE phosphatidylethanolamine, PG phosphatidylglycerol, PI phosphatidylinositol, PS phosphatidylserine, SM sphingomyelin, TG triacylglycerol. **b – g)** Selected analysis of lipid species from untargeted lipidomics classes. Lipid Class specified for each histogram, normalized peak area between AB (grey) and *fam57b mut* (green). Statistical analysis by 2-Way ANOVA, *p ≤ 0.05 **p ≤ 0.01,***p ≤ 0.001, ****p ≤ 0.0001. AB n = 3, *fam57b mut* n= 3, error bars represent SEM. Experiment repeated twice, analysis was similar between two separate runs. Individual MG species between *fam57b het* to AB n.s. Not shown for space: 2-Way ANOVA analysis of TG. Increase in TG (16:0_16:0_16:1) in *fam57b het* compared to AB (p ≤ 0.01).

These data demonstrate that *fam57b* is crucial for regulation of SL and GL classes in the larval zebrafish brain, and that there is a gene dosage-dependent effect. The data in zebrafish brain are quite similar to changes seen in human neurons after FAM57B knockout (**Fig. 4**). These changes affect comparable lipid groups to those altered in 16pdel neurons relative to control, although they are not the same. For example, LPE significantly decreased in the 16pdel neurons (**Fig. 2a**), while the class remained unchanged or significantly increased in the *fam57b* het and null SH-SY5Y cells (**Fig. 4a-c**) and zebrafish brain (**Fig. 6a**,**b**), suggesting additional genes regulate 16pdel lipid metabolites or that these result from other differences between the tissue being compared.

### Changes in plasma membrane and associated proteins in *fam57b mut* and *fam57b het* brains

Ceramide, hexosylceramide and GL species are integral to membrane composition, are differentially distributed across inner and outer leaflets of the plasma membrane and contribute to lipid rafts^47^. Given the changes in these lipids observed in *fam57b mut* brains, we predicted that plasma membrane structure would also be altered. To assess lipid raft organization, fluorophore-conjugated Cholera Toxin subunit B (CT-B), which binds ganglioside GM1 found in lipid rafts^48^, was injected into the hindbrain ventricle of *fam57b mut* and AB zebrafish embryos at 24 hours post fertilization (hpf), when ventricles are accessible for injection^49^ (**Fig. 7a**). Embryos fixed after 1-hour incubation demonstrated a significant increase in punctate GM1 labeling in neural progenitor cells of *fam57b mut* brains compared to AB (**Fig. 7b**). To asses changes in glycerophospholipid species in plasma membranes, we stained with duramycin, a label for membrane PE^50^. Mutant progenitors showed statistically increased punctate PE staining, indicating altered PE localization that could impact membrane architecture (**Fig. 7c**). The duramycin puncta may indicate exosomes or extracellular vesicles containing PE^51^. At 24 hpf, we did not observe changes in cell proliferation or cell death between *fam57b mut* and AB (**Supp. Fig 4**). The data suggest that there is alteration in the plasma membrane of *fam57b mut* brains relative to AB.

**Figure 7.**
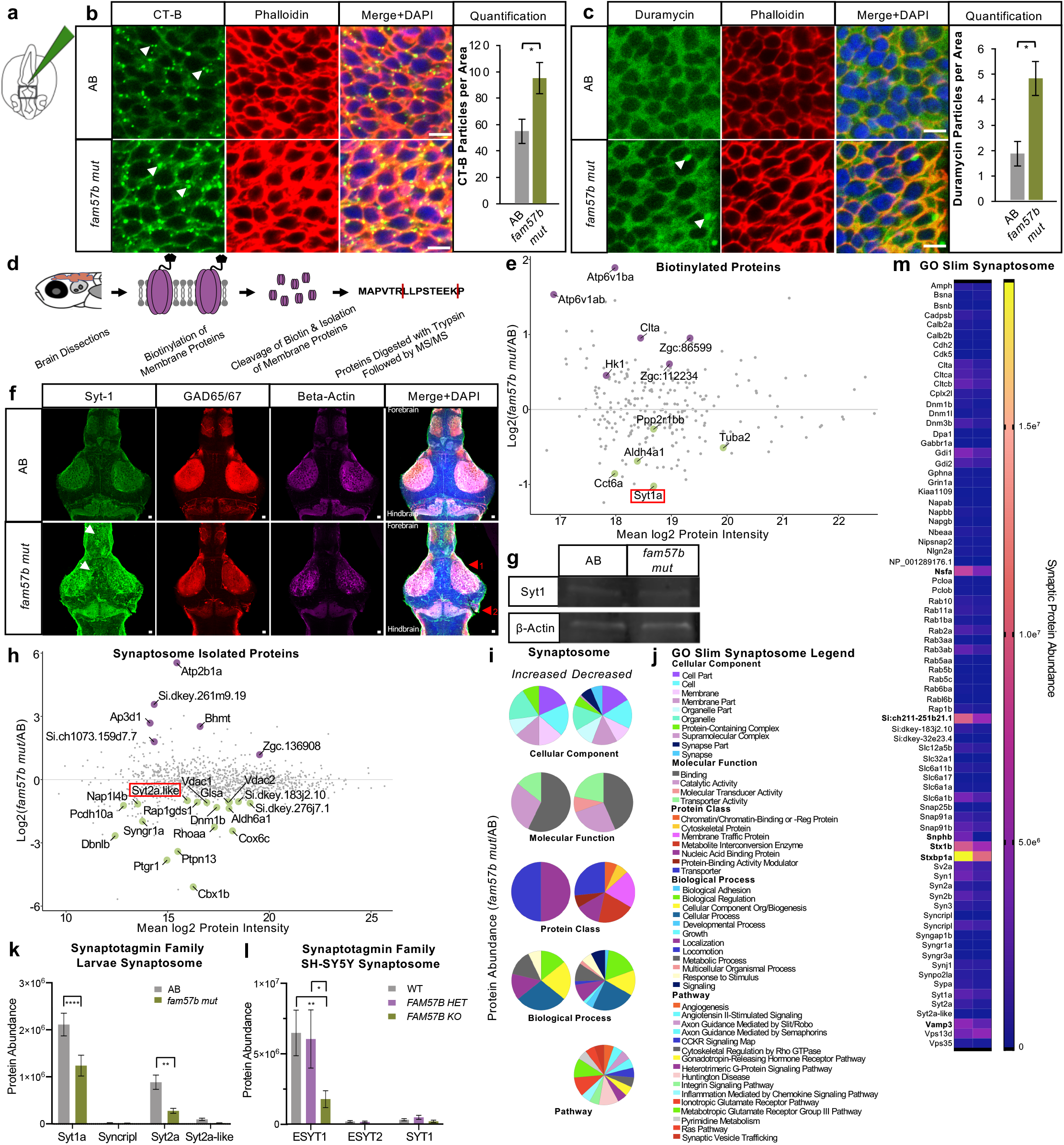
Loss of *fam57b* results in altered plasma membrane architecture early in development and decreased localization of Synaptotagmin family at the synapse later in development. **a)** Schematic of Cholera toxin-B-488 (CT-B) injection into hindbrain ventricle of embryo and flat-mounted midbrain region for imaging at 24 hpf. **b)** Representative embryo midbrain imaging and quantification of CT-B labeling of AB compared to *fam57b mut*. Punctate CT-B labeling (arrows), actin marker phalloidin indicates labelling of CT-B at the plasma membrane, merged with DAPI. Quantification of puncta between WT (grey) and *fam57b mut* (green) CT-B (p ≤ 0.05) T-Test. Scale bar = 5 μm. AB n = 16, *fam57b mut* n = 18. Error bars SEM, statistical analysis by T-test \**p ≤* 0.05. **c)** Representative embryo midbrain imaging and quantification of duramycin-488 labeling of AB compared to *fam57b mut*. Punctate duramycin labeling (arrows), actin marker phalloidin indicates labelling of duramycin at the plasma membrane, merged with DAPI. Quantification of puncta between WT (grey) and *fam57b mut* (green) Duramycin PE staining (p ≤ 0.05) T-Test. Scale bar = 5 μm. AB n = 8, *fam57b mut* n = 8. Error bars SEM, statistical analysis by T-test \**p ≤* 0.05. **d)** Schematic of membrane protein labeling biotinylation assay and processing for MS/MS in 7 dpf larvae brains. **e)** Larvae brain total plasma membrane protein abundance changes between *fam57b mut* relative to AB (Log2 Fold). Statistically significant p ≤ 0.05 −0.0001 proteins labelled, indicating increased (purple) or decreased (green) abundance. Lowest abundance membrane protein Synaptotagmin-1a (red box). n = 6 per genotype. **f)** Representative slice of 7 dpf whole larva brain mount with Sytaptotagmin-1a (green), GAD65/67 (red) and Beta-actin (magenta). Z-stack composite image merged with DAPI. Forebrain and midbrain areas of diffused Syt-1 localization (white arrows). Anatomical differences noted throughout brain, including (1) optic tectum and (2) corpus cerebelli (red arrows). Scale bar = 10 μm. **g)** Representative 7 dpf whole brain western blot indicate no significant change in total Syt-1a protein levels between *fam57b mut* relative to AB. Zebrafish larvae brains pooled (20 per genotype). Syt-1a protein abundance normalized to Beta-Actin loading control, repeated twice. **h)** Larvae brain total isolated synaptosome protein abundance changes between *fam57b mut* relative to AB (Log2 Fold). Statistically significant p ≤ 0.05 −0.0001 proteins labelled, indicating increased (purple) or decreased (green) abundance. Low abundance Synaptotagmin-2a like protein (red box). n = 6 per genotype. **i-j)** Gene ontology analysis of statistically significant larvae synaptosome isolated proteins (h) in *fam57b mut* **i-j)** relative to AB. i) Gene ontology pie graphs of increased and decreased protein groups of cellular components, molecular function, protein classes, biological processes and pathways. j) Gene ontology figure legend. **k)** Analysis of Synaptotagmin family members from larvae isolated synaptosomes. Significantly decreased protein abundance of Syt1a and Syt2a by 2-Way ANOVA, **p ≤ 0.01, ****p ≤ 0.0001. **l)** Analysis of Synaptotagmin family members from differentiated SH-SY5Y isolated synaptosomes between all 3 genotypes. Significantly decreased protein abundance of elongated ESYT1 by 2-Way ANOVA, *p ≤ 0.05, **p ≤ 0.01. Error bars SEM. **m)** Analysis of synaptic markers from larvae isolated synaptosomes. Significantly decreased protein abundance (bolded) of vesicle regulation machinery and glutamate receptor activity by 2-Way ANOVA, **p ≤ 0.01, ****p ≤ 0.0001.

The changes in membranes of neural progenitor cells suggested that membrane protein localization may also be altered. We therefore examined localization of membrane by biotinylation analysis. Freshly dissected larval brains from *fam57b mut* or AB at 7 dpf were incubated with membrane impermeable biotin. Surface proteins were affinity-purified and quantified by MS/MS (**Fig. 7d)**. MS/MS analysis indicated that membrane-associated protein cohorts were similar between *fam57b mut* and AB brains (**Fig. 7e**), however, a small group of proteins showed altered abundance. In *fam57b mut* brains, the protein whose levels most significantly decreased (2-fold) relative to AB was Synaptotagmin-1a (Syt1) (**Fig. 7e**). Syt1a, homologous to human SYT1, is a vesicle membrane protein that acts as a calcium sensor and regulates synaptic and endocrine vesicle exocytosis^52–56^. Syt1 protein domains interact with the lipid bilayer, including GL PS that are altered in *fam57b mut* and *fam57b het* (**Fig. 5a**,**b**). Mammalian Syt1 can modify PS, and is able to alter curvature strain on the membrane^57^.

To investigate the decreased membrane abundance of Syt1 in the biotinylation assay, we performed immunostaining on 7 dpf larvae brains (**Fig. 7f**). Whole larval brains were cleared and tertiary structure was protected using SHIELD protocols. Slice imaging of dorsal brain view showed that Syt1 protein was largely confined to projections of neurons throughout the AB brain, both GABAergic (GAD65/67) and non-GABAergic, while *fam57b mut* brains showed ectopic expression throughout the brain (**Fig. 7f**). By western blot, we found no change in total brain Syt1 between *fam57b mut* and AB, suggesting that immunostaining demonstrates Syt1a mislocalization (**Fig. 7g**). Imaging also revealed anatomical changes in the larval *fam57b mut* brain, including tectum and corpus cerebelli (**Fig. 7f**).

Together, these data indicate that in the brain, relative to wildtype, *fam57b mut* animals show changes in lipids, membrane structure and membrane protein association, including the synaptic regulator Syt1 and others functioning at synapses. These data indicate that *fam57b* is required for membrane structure and neuronal architecture.

### Pre-and post-synaptic proteins depleted after loss of *FAM57B*

To examine the implications of Syt1 mis-localization on synaptic composition, we isolated synaptosomes from fresh brains of *fam57b mut* or AB larvae. Proteomic profiling indicated a significantly increased group of proteins, and a larger significantly decreased group of proteins in *fam57b mut* compared to AB (**Fig. 7h**). Interestingly, we observed a decrease of the Synaptotagmin family member Syt2a-like protein, similar to human SYT2, with relatively analogous function to SYT1. Gene ontology (GO-Slim and Panther Protein Class ontology) defined synaptic protein groups only found in the decreased synaptosome protein group (**Fig. 7i, 7j**). Annotations in the decreased group included synapse and synapse part components, cytoskeletal and membrane traffic proteins, biological adhesion, development and signaling, and numerous implicated pathways including synaptic vesicle trafficking. These results indicate that synaptic protein levels were decreased in synaptosomes from *fam57b mut* larval brain synapses relative to AB.

We separately examined levels of synaptotagmin family members in the brain synaptosome profiles and found decreased Syt1 and Syt2a protein levels in *fam57b mut* compared to AB synaptosomes (**Fig. 7k**). This interesting and significant association between FAM57B regulation and Synaptotagmin expression (**Fig. 7e**,**f**,**h**) led us to probe further into analysis from human SH-SY5Y isolated synaptosomes. We found a significant decrease in elongated SYT1 (ESYT1) in *FAM57B KO* compared to *FAM57B HET* and WT (**Fig. 7l**), a calcium activated synaptic protein found to bind GLs^58^. We then characterized hallmark proteins that function at the synapse from brain synaptosomes, including synaptic vesicle fusion and tethering proteins. Bayés *et al*. previously examined complexity of the adult zebrafish synapse proteome relative to adult mouse synapse proteome^59^. As expected, not all synaptic proteins were detected at this immature stage of zebrafish development compared to the adult brain. Enrichment of synaptic vesicle proteins including Syntaxins, Slc neurotransmitter transporters, SNAPs, Stx/Vps, Synaptotagmins and membrane budding proteins including Dynamins and Rabs verify synaptosome isolation and give new data regarding neuronal maturation in the zebrafish larval brain (**Fig. 7j**). Comparison of synaptic protein profiles between genotypes demonstrated decreased vesicle fusion and transport protein Nsfa, ligand-gated ion channel Si:ch211-251b21.1, and SNARE complex proteins Snphb, Stx1b, Stxbp1a and Vamp3 (**Fig. 7j**). Together, these data show that synaptic proteins essential for vesicle docking, exo-and endocytosis, including synaptotagmin family members, are diminished in synaptosomes isolated from *fam57b mut* brains relative to AB, suggesting Fam57b is essential for synapse integrity.

### Depressed spontaneous electrical activity and response to stimuli in *fam57b* mutants

To understand how changes in *fam57b* gene dosage impact neuronal activity, we tested brain activity by electrophysiological analysis. We previously described a noninvasive electrophysiology technique that can be used in live larvae to measure spontaneous activity in the brain and spinal cord^43^. Using a multielectrode array (MEA), we measured local field potential (LFP) parameters and relative coordinated (network) activity in the brain of 7 dpf larvae (**Figs. 8a-c**). Larva were individually immersed in precooled 1.5% low-melt agarose in E3 solution and mounted in a 64-electrode containing well. We measured spontaneous brain activity over a 10-minute period, comparing *fam57b mut* to AB controls. Only electrodes in contact with the larval head were analyzed, ∼6 to 8 electrodes, whose signal was pooled. *fam57b mut* larvae had slightly smaller heads than AB at 7 dpf (**Supp Fig. 5**), however these changes do not impact electrophysiological studies. Overall, *fam57b mut* spontaneous brain activity was severely diminished relative to ABs. This included significant decrease in number of LFPs, mean LFP rate, and inter-LFP-interval coefficient of variation measurements, indicating DO decreased spontaneous brain activity with reduced kinetics (**Fig. 8a**). However, the decreased coefficient of variation in the *fam57b mut* suggests LFP interval distributions are detected at a more regular rate. Measuring electrographic bursts, at least 5 LFPs per 100 ms, we were unable to detect any burst activity under these settings in the *fam57b mut*, while bursts were detected in ABs. To increase sensitivity for detection of burst activity, we lowered the detection parameters to at least 3 LFPs per 200 ms (right column), and found decreased electrographic burst duration, number of LFPs per burst, burst frequency and percentage in *fam57b mut* relative to AB (**Fig. 8a**). In addition, we examined relative network activity, as defined by at least 3 LFPs detected simultaneously between a minimum of two electrodes. Relative network activity was also significantly decreased in *fam57b mut* compared to AB. Synchrony index of bursts did not change between the two genotypes, indicating coordination of network activity did not differ. While LFP waveforms could not be quantified due to small distance variations when mounting individual larva, we observed smaller relative waveforms in *fam57b mut* compared to AB (**Fig.8c**), consistent with overall decreased brain activity in *fam57b mut* larvae. A representative raster plot of LFP activity in the head region over the 10-minute recording period illustrates the relative decrease in LFP propagation, burst and network detection measured (**Fig. 8d**). A representative image of a mounted larva immersed in agarose on a 12-well 64 electrode plate is shown in **Fig. 8e**. These data demonstrate severely diminished spontaneous brain activity in *fam57b mut* relative to AB wildtype larvae, and highlight a role for Fam57b in regulating brain function.

**Figure 8.**
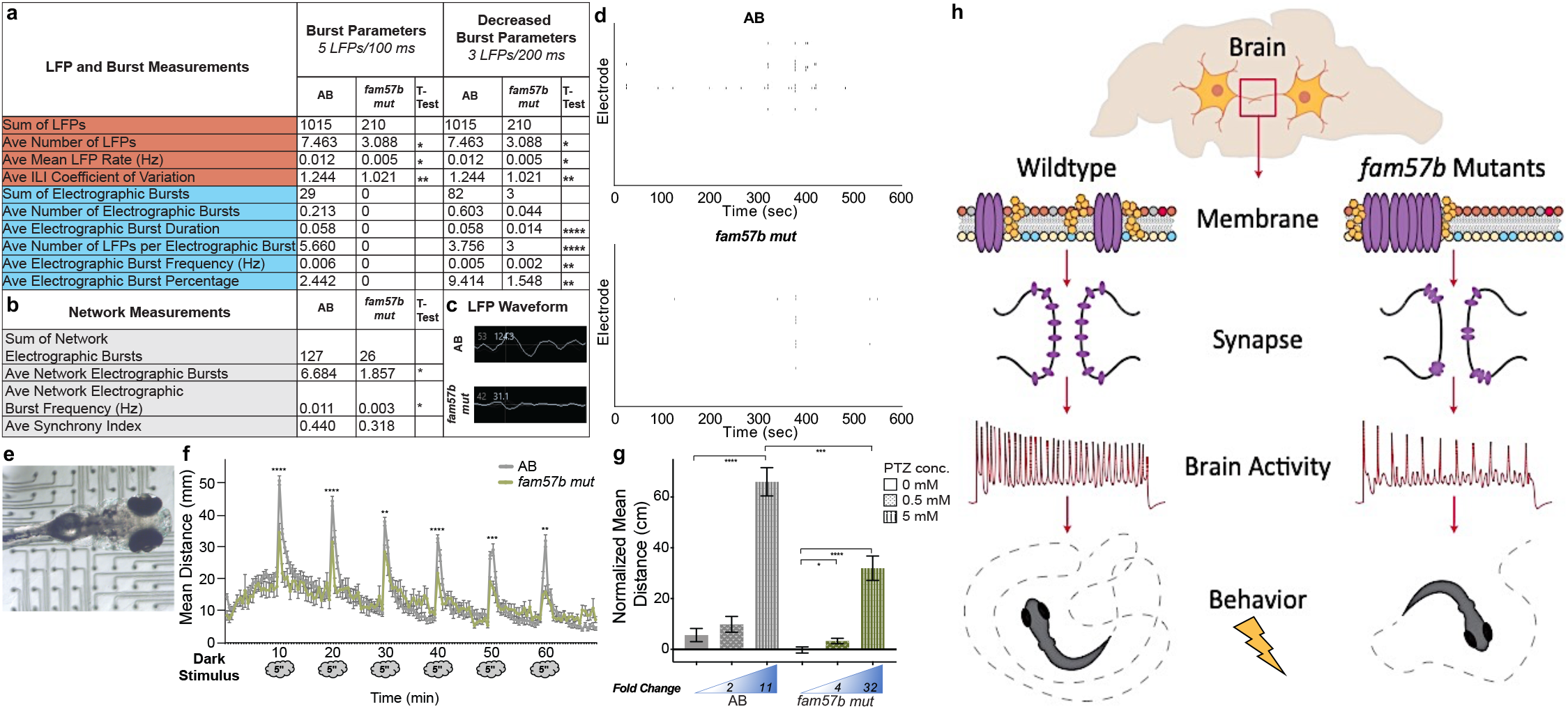
Decreased spontaneous brain activity and diminished behavioral response after stimuli presentation in *fam57b* mutants. **a-b)** Local field potential (LFP) recordings in unanesthetized live larvae at 7dpf. Brain localized LFP recordings were pooled for each larva. **a)** Decreased average number, mean rate and inter-LFP interval (ILI) coefficient of variation of LFP in *fam57b mut* compared to AB (orange). No electrographic burst activity was identified in *fam57b mut* at standard 5 LFPs/100 ms. Decreased electrographic burst parameters, including duration, number of LFPs per burst, frequency and percentage at 3 LFPs/200ms (blue). **b)** Decreased average electrographic burst network activity and frequency, defined as a minimum or 3 electrographic bursts between 2 electrodes simultaneously, in *fam57b mut* compared to AB (grey). AB n = 21, *fam57b mut* n = 24 over 6 experiments. Statistical significance by unpaired T-test, *p ≤0.05, **p ≤ 0.01,*** p ≤ 0.001,**** p ≤ 0.0001. **c)** Representative LFP waveform in brain region, indicating smaller relative waveform in *fam57b mut* compared to AB. **d)** Representative LFP raster plot over experimental time frame, indicating less overall activity in *fam57b mut* compared to AB. **e)**Representative image of 7 dpf immersed in cooled agarose in contact with electrodes on 12-well CytoView MEA plate. **f)** Startle response behavioral assay. Light source was removed for 5 secs at 10 min intervals. Mean distance reported from tracked movement during 70 min assay. Decreased light startle response identified in *fam57b mut* compared to AB. Statistical analysis of each startle response by T-test **p ≤ 0.01, *** p ≤ 0.001, **** p ≤ 0.0001. Error bars SEM. No overall significant change in movement outside of the startle identified. AB n = 125, *fam57b mut* n = 33 over 5 experiments. **g)** Seizure response behavioral assay. Normalized mean distance from tracked movement after absence or presence of pentylenetetrazol (PTZ) 0.5 mM and 5 mM. Significantly increased seizure-induced movement observed at 5mM in AB, while increased movement observed at 0.5 and 5 mM PTZ in *fam57b mut*. Diminished overall seizure-induced movement at 5 mM in *fam57b mut* compared to AB. Relative fold change compared to absence of PTZ indicated below histogram. Statistical analysis of each condition by T-test *p≤ 0.05, *** p ≤ 0.001, **** p ≤ 0.0001. Error bars SEM. AB n = 166 (0 mM), 92 (0.5 mM), 150 (5 mM), *fam57b mut* n = 91 (0 mM), 70 (0.5 mM), 56 (5mM) over 6 experiments. **h)** Model proposing role of Fam57b activity in the brain. Loss of function in *fam57b* mutants indicate significant changes in plasma membrane lipid groups alter architecture of plasma membrane early the developing brain. Architectural changes indicated by increased lipid raft abundance and aggregation. Altered plasma membrane homeostasis results in mis-localization of synaptic proteins, including synaptotagmins, after maturation. Decreased spontaneous brain and network activity suggests diminished synaptic function and developed circuits. As evidence has suggested spontaneous network activity shapes synaptic development, this cycles back to declined neuronal maturation and circuity. Molecular changes to synaptic function and decreased spontaneous brain activity translate to altered behavioral response after stimuli presentation.

After identifying significantly diminished spontaneous brain activity in *fam57b mut* larvae, we examined correlations with behavioral activity. We first tested light-responsive sensorimotor startle behavior. Startle response, as indicated by distance traveled, was measured over a 70-min time-frame with light extinguished every 10 minutes for 5 seconds^43^. The startle response window was in total 30 seconds, including the stimulus. We found a considerable decrease in response to each light stimulus in *fam57b mut* compared to AB (**Fig. 8f**). However, movement measured previous to startle (first 10 min) and relative habituation after startle cue did not overall differ between the genotypes, indicating there is no alteration in movement outside of the light stimulus, and no visual deficit in *fam57b mut* larvae.

To examine brain specific activity, we investigated seizure susceptibility. Seizures are prevalent in individuals affected with 16pdel syndrome, and may result from processes involving several neurotransmitter systems, including glutamatergic, cholinergic and GABAergic^60^. To measure seizure propensity, larvae were immersed in pentylenetetrazol (PTZ), a GABA_A_ antagonist, well characterized for use in zebrafish^61^. After a 10-minute baseline movement recording, two different concentrations of PTZ, or E3 media only control, were applied to the individual well of each larva and recorded over 10 minutes. There was no significant change in normalized movement in the absence of stimulus after addition of E3 (0 mM PTZ) control between the *fam57b mut* and AB (**Fig. 8g**). Increasing the PTZ dose increased normalized distance traveled for both genotypes, but the increase was not significant for AB between 0 and 0.5 mM PTZ as observed in *fam57b mut*. At 5 mM, we observed significantly less distance traveled in the *fam57b mut* compared to AB. However, the relative fold change between 0 and 5 mM was much higher in the *fam57b mut* (roughly 32-fold) compared to AB (roughly 11-fold). These analyses indicate neuronal specific changes after loss of *fam57b mut*, however, FAM57B is also expressed in muscles. Staining the neuromuscular junction, we did not observe any overt differences between *fam57b mut* and AB (**Supp. Fig. 6**). Together, altered brain activity, response to multiple stimuli including GABA_A_ antagonist, and gross anatomical differences including the corpus cerebelli (**Fig. 7f**), suggest significant changes to GABAergic network activity in the developing *fam57b mut* brain.

In sum, there are significant behavioral changes in zebrafish larvae after fam57b loss of function. *fam mut* larvae move similar to AB over time without and after recovery of a stimulus. However, with a stimulus – either dark or PTZ application, there is altered behavioral responsiveness. These findings are consistent with changes in brain activity in *fam57b mut* relative to AB controls.

## Discussion

This study has uncovered alterations in neurons derived from people affected with 16pdel Syndrome, that in a neuronal cell line and in the zebrafish model occur concomitantly with changes in cellular lipid composition, membrane architecture, membrane associated proteins, synaptic activity, brain activity and behavior. We do not know whether these outcomes are linked, but it is plausible that FAM57B acts through SL and GL regulation that is the starting point for a cascade of effects in loss of function or mutation. Dysregulated lipid metabolism exerts a multifaceted effect on neurons. Increased lipid energy consumption escalates oxidative stress, exacerbating inflammation, mitochondrial and metabolic dysfunction and excitotoxicity. The structure of individual lipid classes affects their intracellular localization, impacting cytoskeleton and lipid raft composition, disrupting signaling processes crucial in regulating neurotransmitter synthesis and release, cytoskeletal integrity, myelination and intracellular transport^62,63^. Well documented myelination dysfunction is prevalent in 16pdel syndrome, with a widespread pattern of aberrant white matter microstructure associated with impaired cognition characteristic of this CNV^38^. Abnormal cholesterol metabolism has been observed in patients with the ASD Asperger syndrome and other non-syndromic ASDs, suggesting a correlation between lipid raft formation and ASD^64,65^. Interestingly, beyond *FAM57B*, multiple genes in the 16p11.2 interval encode enzymes with predicted roles in metabolic processing or interconversion including *ALDOA, CDIPT, GDPD3, BOLA2, SULT1A3, SULT1A4* and *YPEL3*^*66,67*^.

The above studies provide evidence for FAM57B enzymatic activity and its functional role in the brain. In cells from individuals with 16pdel Syndrome, we found that iPSC differentiated cortical neurons display increased excitability when assessing network activity, and uncovered a sex-linked difference between 16pdel males and females. We determined that 16pdel neurons have a significantly altered lipidome, indicating changes in metabolism of ceramides, SLs and GLs that are critical for function between the ER, mitochondria and the plasma membrane. Through ceramide synthase activity studies, we identified that FAM57B is in fact a ceramide synthase modulator. Lipidomic profiling uncovered altered SLs and GLs integral to the lipid membrane in *FAM57B* mutants, with similar lipid group disruption in 16pdel iPSC differentiated patient neurons and neuronal SH-SY5Y cells. These results support our findings of interaction between FAM57B with CerS2 and CerS6, and indicate FAM57B is necessary for proper neuronal development. Early in brain development, *fam57b* mutants displayed altered plasma membrane architecture. As neurons matured, we determined synaptic proteins were significantly diminished in mutants. Diminished spontaneous brain activity and altered behavioral response after stimulation supported major changes in synaptic composition. These studies suggest Fam57b is essential early in development of the brain, and loss of *fam57b* leads to cascading events starting with changes in plasma membrane architecture followed by disturbance in protein organization at the membrane and detriments to basic neuronal function. Together, we propose a model whereby Fam57b functions to maintain normal plasma membrane physiology, necessary for proper formation and function of neurons (**Fig. 8h**).

Syt1 acts as an essential determinant of synaptic vesicle endocytosis by delaying the kinetics of vesicle retrieval in response to increasing Ca^2+^ levels^68^. Interestingly, ESYT1 may play a role in cellular lipid transport of PC, PE, PI, and translocates to sites of contact between the presynaptic endoplasmic reticulum and the cell membrane in response to increased cytosolic calcium levels^58^. Neurotransmission is decreased in *Esyt D*.*melanogaster* mutants, with a proposed role in synapse extension, highlighting the essential homeostasis of lipids at the synapse^69^. SYT1-associated neurodevelopmental disorder, also known as Baker-Gordon Syndrome, is an autosomal dominant heterozygous mutation of *SYT1*. Mutations are found to disrupt SYT1 function causing a reduction in neurotransmitter release, and is associated with infantile hypotonia, congenital ophthalmic abnormalities, hyperkinetic movement disorder, stereotypical motor behavior, intellectual disability, sleep disturbance and episodic agitation^70^. While a dominant mutation, the heterozygous loss of SYT1 provides insight into the depressed electrophysiological and behavioral responses, and probes further investigation of SNARE complex protein abundance and function in 16pdel Syndrome. This association is particularly important with our previous findings of a genetic interaction between *fam57ba* and calcium-sensitive exocytosis regulator *doc2a*, another gene within the 16p11.2 interval, with double heterozygotes showing hyperactivity and increased seizure propensity^16^.

A recent study analyzed the largest exome sequencing study of ASD to date, with 35,584 total samples and 11,986 with ASD^71^. This study identified 102 high risk genes tightly associated with ASD. In the analysis, a rare G:A mutation was discovered in *FAM57B*, located in the 5’UTR coding exon of one *FAM57B* transcript, and resides in the promoter/enhancer region of the 5 other transcripts. This synonymous mutation is predicted to create binding sites for several transcription factors and could potentially impact predicted enhancer activity in neurons, affecting transcription, mRNA stability and/or splicing (prediction with information from dbSNP, JASPAR, GTEx). This study indicates a potential *FAM57B* mutation associated with ASD outside of CNV 16pdel, and encourages further evaluation of this enzyme to brain development and function.

Our approach focused on lipid metabolites in understanding neurodevelopmental and mental health disorders. While *FAM57B* haploinsufficiency alone cannot account for the multitude of disrupted biochemical and cellular properties in 16pdel patient neurons, our analysis highlights disrupted metabolism as tightly correlated to the syndrome. By combining lipidomics with proteomics, biochemistry, cellular biology, and neuronal activity to understand function of FAM57B, we can better link the landscape of changes to synaptic homeostasis.

## Materials & Methods

### Fish lines and maintenance

Adult zebrafish of the wildtype AB strain were maintained at 28°C on 12h/12h light/dark cycle. Embryos were obtained from natural spawning and staged as previously described by Kimmel *et al*.^72^. Due to the polygenic nature of sex determination and timing of gonadal development in zebrafish, we are unable to determine the sexes of the embryos and larvae for our assays. However, because our assays utilized large numbers of embryos and larvae, both sexes should be adequately represented. Embryos were obtained from separate crosses of *fam57b mut* mutant fish. *fam57b het* fish were generated by crossing *fam57b mut* to AB fish.

*fam57ba*^*-/-*^ mutants were injected with *fam57bb* targeted sgRNA at 1 −4 cell stage, previously described in McCammon *et al*. CRISPR/Cas9 induced mutation resulted in 17 bp deletion and early stop codon. Experiments were performed after 4 generations of crosses with AB controls. fam57bb 5’ to 3’ TAGGTGATGTCCTGGCAGGAAG fam57bb 3’ to 5’ AAACCTTCCTGCCAGGACATCA For genotyping, PCR amplified region of in/del. PCR was digested with EarI restriction enzyme, with homozygous mutation detected by loss of EarI restriction site. *fam57b mut* line was outcrossed with AB periodically to avoid chromosomal abnormalities.

### Induced Pluripotent Human Stem Cells

Unaffected control, 599 and 657, iPSC lines were a generous gift from Rudolf Jaenisch, originally obtained as fibroblasts from Corriell Institute Biobank. iPSC of 16p11.2 deletion carriers were obtained from Simons Variation in Individuals Project^22^. Cell line corresponds to subjects with abbreviated ID from RUCDR. All iPSCs were tested for negative mycoplasm and normal karyotype. Cytogenetic analysis was performed on twenty G-banded metaphase cells at Cell Line Genetics. All experiments involving cells from human donors were performed in compliance with established IRB protocols at the Whitehead Institute.

*iPSCs* -Cells were cultured on plates coated with Matrigel (Corning) in mTeSR media (STEMCELL Technologies) with pen/strep. Y27632 was added to cells prior to passaging, then passaged with ReLeSR (STEMCELL Technologies). Cells were maintained at 37°C with 5% O_2_.

*Generation of Cortical Neurons* -iPSCs were differentiated into neural progenitor cells (NPCs) by FGF exchange. FGF was slowly removed by exchanging H9 media (DMEM/F12/HEPES, Knockout SR (Gibco), NEAA, GlutaMAX (Gibco), Beta-Mercaptaethanol, pen/strep and Human FGF (Peprotech) with -FGF media (DMEM/F12/HEPES, Neurobasal (Thermo), N2 (Gibco), Gem21 (Gibco), non-essential amino acids (NEAA), GlutaMAX (Gibco), pen/strep, D-Glucose and NaCl) plus dorsomorphin (Tocris) every day over 2-3 weeks. When rosettes were present, cells were incubated with Y27632 and passaged with Accutase (STEMCELL Technologies). Cells were expanded and passaged at least 3 times until homogeneous NPC culture. NPCs were passaged on poly-D-lysine and laminin coated plates for cortical neuron differentiation. NPCs were incubated with Neuronal Differentiation media (Neurobasal, GlutaMAX, NEAA, D(+) Glucose (Sigma), Gem21, Culture One (Gibco), BDNF and GDNF (Peprotech), pen/strep) for 1 month, changing media every 2 to 3 days. Cells were maintained at 37°C under normoxic conditions.

### SH-SY5Y Neuroblastoma cell line

SH-SY5Y cells, originally from ATCC, were a kind gift from David Bartel, Whitehead Institute for Biomedical Research. Cells were maintained in EMEM, F12 media supplemented with fetal bovine serum and pen/strep in a 37°C incubator with 5% CO_2_. Differentiation of cells to neuronal model were induced with media containing Neurobasal, Gem21, GlutaMAX, All-trans-retinoic acid (Sigma) and pen/strep, for 4 days in dark to prevent retinoic acid degradation from light exposure^33^.

CRISPR sgRNA designs were identified from Target Guide Sequence Cloning Protocol, Zhang lab, with sequence overlapping the TLC domain of FAM57B^73^. 10 targeted guides to FAM57B sequence were individually transformed in pLC OPTI-Stuffer plasmid, a kind gift from David Sabatini, Whitehead Institute, and lentivirus was grown in HEK293 cells. Generation of CRISPR/Cas9 induced mutations via lentiviral transduction was performed according to Wang *et al*. protocols^74,75^. After puromycin selection, cells were gently triturated and diluted to approximate 1 cell per well in 96 well plate. Wildtype cells were simultaneously single cell diluted and sorted to serve as additional control for experiments. Incorporation of mutation was determined by Next Generation Sequencing.

FAM57B Homozygote deletion (KO)

sgFAM57B 5’ to 3’ -GGTGCTCCACCATGCCGCCA

Mutation resulted in frameshift with 111 and 121 bp deletion on either strand, resulting in early stop codon.

FAM57B Heterozygote deletion (HET)

sgFAM57B 5’ to 3’ -GGGCACAGCAAATTGCGTGT

Mutation resulted in frameshift with 20 bp deletion on one strand, resulting in early stop codon.

Adeno-Associated Virus Integration Site 1 (AAVS1) targeted control sgAAVS1 5’ to 3’ -CACCGGGGCCACTAGGGACAGGAT

Mutation resulted in frameshift and 51 bp and 1 bp deletion on either stand, resulting in early stop codon. The AAVS1 served as a control for all SH-SY5Y experiments. WT was compared to AAVS1 to determine confidence of statistical significance when compared to *FAM57B HET* and *KO*.

### HEK293 Cell Culture and Co-Immunoprecipitation

HEK293T cells were cultured in Dulbecco’s modified Eagle’s medium supplemented with 10% fetal bovine serum, 100 IU/ml penicillin, 100 µg/ml streptomycin, and 110 µg/ml sodium pyruvate. Transfections were performed with the polyethyleneimine reagent using 8 µg of plasmid per 10 cm culture dish for 36–48 h; medium was exchanged after 6 hours. pcDNA3.1 was used as a control. DYKDDDDK (Flag)-tagged human FAM57B plasmid was purchased from Genscript. Hemagglutinin (HA)-tagged human CerS plasmids were generated as described^76^.

Co-immunoprecipitation was performed using cells transfected with a variety of plasmids. HA-tagged CerSs were used to confirm non-specific binding to Flag affinity resins. Cells were washed twice with cold PBS and lysed in lysis buffer (20 mM Tris (pH 7.4), 150 mM NaCl, 1 mM EDTA, 1% NP-40, 5% glycerol and protease inhibitor (Sigma-Aldrich)). Lysates were incubated on ice for 10-15 min. Protein was determined using the BCA reagent. FLAG-tagged human FAM57B, using an anti-FLAG affinity resin (Genscript). Lysates were incubated with 40 µl of beads overnight at 4°C with rotation. The resin was washed three times in 1 ml of lysis buffer at 4 °C with rotation. Proteins were eluted using 4X SDS sample buffer (BioRad). Eluted proteins were analyzed by Western blotting for detection of HA-tagged interacting proteins.

### Ceramide Synthase Assays

Cell homogenates were prepared in 20 mM HEPES-KOH, pH 7.2, 25 mM KCl, 250 mM sucrose, and 2 mM MgCl2 containing a protease inhibitor mixture. Protein was determined using the BCA reagent (Thermo Fisher Scientific). Samples were incubated with 15 µM NBD-sphinganine (Avanti Polar Lipids), 20 µM defatted BSA (Sigma-Aldrich), and 50 µM fatty acyl-CoA (Avanti Polar Lipids) in a 20 µl reaction volume. CerS (40 µg protein, 25 min reaction time) was assayed using C24.1-CoA and Cer5/6 (5 µg protein, 5 min reaction time) assayed using C16-CoA. Reactions were terminated by chloroform/methanol (1:2, v/v) and lipids extracted. Lipids were dried under N2, resuspended in chloroform/methanol (9:1, v/v), and separated by TLC (Merck) using chloroform/methanol, 2M NH4OH (40:10:1, v/v/v) as the developing solvent. NBD-labeled lipids were visualized using an Amersham Typhoon5 imager and quantified by ImageQuantTL (GE Healthcare, Chalfont St Giles, UK). All solvents were of analytical grade and were purchased from Bio-Lab (Jerusalem, Israel).

### Sample Collection for Lipidomics

#### iPSC Differentiated Neurons

2 × 10^6^ NPCs were plated in 6 well plate, at least 3 wells per genotype. Differentiation to cortical neurons was performed as stated above. Cells were washed with phosphate buffered saline solution (PBS). Cells were scraped in LC grade methanol and homogenized in eppendorf tube containing water and LC grade chloroform with pestle mixer, followed by vortexing for 10 minutes at 4°C. Lipids were separated by centrifuging top speed at 4°C. This was repeated three times with all samples run together in positive ion mode for lipidomic analysis.

#### Zebrafish

At 7 dpf, larvae were deeply anesthetized with tricaine. Brains with surrounding epidermal layer were dissected and flash frozen on dry ice. Collections were pooled at 20 brains per sample. Brains were homogenized in an eppendorf tube with a pestle mixer in LC grade methanol, LC grade chloroform and water, followed by vortexing for 10 minutes at 4°C. Lipids were separated by centrifuging top speed at 4°C. Brains were collected and stored in −80°C over many dissections to acquire adequate tissue for analysis. This was repeated twice per genotype, with the *fam57b mut* and AB cohort run at different times (different normalization) while the *fam57b het* and AB cohort were run at the same time.

#### Differentiated SH-SY5Y Cells

1 × 10^6^ cells were plated per well in 6 well plate, 3 wells per genotype. SH cells were differentiated over 4 days in media containing retinoic acid. Cells were washed with PBS. Cells were scraped in LC grade methanol and homogenized in eppendorf tube containing water and LC grade chloroform with pestle mixer, followed by vortexing for 10 minutes at 4°C. Lipids were separated by centrifuging top speed at 4°C. This was repeated twice with all samples run together.

### Untargeted Lipidomics

Lipids were separated on an Ascentis Express C18 2.1 × 150 mm 2.7 um column (Sigma-Aldrich) connected to a Vanquish Horizon UPLC system and an ID-X tribrid mass spectrometer (Thermo Fisher Scientific) equipped with a heated electrospray ionization (HESI) probe. External mass calibration was performed using the standard calibration mixture every seven days. Dried lipid extracts were reconstituted in 50 uL 65:30:5 acetonitrile: isopropanol: water (v/v/v). Typically, 2 uL of sample were injected onto the column, with separate injections for positive and negative ionization modes. Mobile phase A in the chromatographic method consisted of 60:40 water: acetonitrile with 10 mM ammonium formate and 0.1% formic acid, and mobile phase B consisted of 90:10 isopropanol: acetonitrile, with 10 mM ammonium formate and 0.1% formic acid. The chromatographic gradient was adapted from Hu *et al*. (2008)^77^ and Bird *et al*. (2011)^78^. Briefly, the elution was performed with a gradient of 40 min; during 0−1.5 min isocratic elution with 32% B; from 1.5 to 4 min increase to 45% B, from 4 to 5 min increase to 52% B, from 5 to 8 min to 58% B, from 8 to 11 min to 66% B, from 11 to 14 min to 70% B, from 14 to 18 min to 75% B, from 18 to 21 min to 97% B, during 21 to 35 min 97% B is maintained; from 35−35.1 min solvent B was decreased to 32% and then maintained for another 4.9 min for column re-equilibration. The flow rate was set to 0.260 mL/min. The column oven and autosampler were held at 55°C and 15°C, respectively. The mass spectrometer parameters were as follows: The spray voltage was set to 3.25 kV in positive mode and 3.0 kV in negative mode, and the heated capillary and the HESI were held at 300°C and 375°C, respectively. The S-lens RF level was set to 45, and the sheath and auxiliary gas were set to 40 and 10 units, respectively. These conditions were held constant for both positive and negative ionization mode acquisitions. The mass spectrometer was operated in full-scan-ddMS/MS mode with an orbitrap resolution of 120,000 (MS1) and 30,000 (MS/MS). Internal calibration using Easy IC was enabled. Quadrupole isolation was enabled, the AGC target was 1×10^5^, the maximum injection time was 50 msec, and the scan range was m/z = 200-2000. For data-dependent MS/MS, the cycle time as 1.5 sec, the isolation window was 1, and an intensity threshold of 1×10^3^ was used. HCD fragmentation was achieved using a step-wise collision energy of 15, 25, and 35 units, and detected in the orbitrap with an AGC target of 5×10^4^ and a maximum injection time of 54 msec. Isotopic exclusion was on, a dynamic exclusion window of 2.5 sec was used, and an exclusion list was generated using a solvent bank.

High-throughput annotation and relative quantification of lipids was performed using LipidSearch v4.2.21 (ThermoFisher Scientific/ Mitsui Knowledge Industries) using the HCD database^79,80^. LipidSearch matches MS/MS data in the experimental data with spectral data in the HCD database. Precursor ion tolerance was set to 5 ppm, product ion tolerance was set to 10 ppm. LipidSearch nomenclature uses underscores to separate the fatty acyl chains to indicate the lack of *sn* positional information. In cases where there is insufficient MS/MS data to identify all acyl chains, only the sum of the chains is displayed. Following the peak search, positive and negative mode data were aligned together where possible and raw peak areas for all annotated lipids were exported to Microsoft Excel and filtered according to the following predetermined quality control criteria: Rej (“Reject” parameter calculated by LipidSearch) equal to 0; PQ (“Peak Quality” parameter calculated by LipidSearch software) greater than 0.75; CV (standard deviation/ mean peak area across triplicate injections of a represented (pooled) biological sample) below 0.4; R (linear correlation across a three-point dilution series of the representative (pooled) biological sample) greater than 0.9. Typically, ∼70% of annotated lipids passed all four quality control criteria. Redundant lipid ions (those with identical retention times and multiple adducts) were removed such that only one lipid ion per species/ per unique retention time is reported in merged alignments. For data where positive and negative mode data were aligned separately some redundancies may still exist. Raw peak areas of the filtered lipids were normalized to total lipid signal (positive or negative ionization mode) in each sample to control for sample loading. Data presented are shown as Log_2_FC compared to wildtype/control samples. Statistics were performed in Prism, with each run analyzed separately.

### Zebrafish brain staining and imaging

For 24 hours post-fertilization staining, embryos were deeply anesthetized in tricaine after being dechorionated. Embryos were places into wells in 1% Agarose dishes. 1 ng Cholera Toxin subunit B (CT-B) (Recombinant Alexa Fluor 488 conjugate, Life Technologies) was injected into the hindbrain ventricle^81^. Embryos were washed with E3 and incubated for 1 hour to allow CT-B binding. Embryos were then fixed in fresh 4% PFA in phosphate buffered solution (PBS) overnight at 4°C. Embryos were washed in PBS + Tween-20 (PBT) and incubated with 555-Phalloidin for 1 hour. Alternatively, embryos were incubated with Duramycin-fluro conjugate (Molecular Targeting Technologies) PE stain for 45 min. Embryos were washed in PBT and mounted in DAPI Antifade (Thermo) overnight. Imaging was performed on an inverted Zeiss LSM700 Laser Scanning Confocal and processed on Fiji (ImageJ). CT-B and Duramycin images were processed on ImageJ to measure relative puncta from staining. Particles were measured after drawing a size circle in each hemisphere comparing AB to *fam57b mut* embryos. Threshold was set to intermodes to assume for bimodal histogram, particle size set between 0 – 2 μm^2^.

For 7 dpf larvae, the following protocol was adapted from the mouse protocol provided by LifeCanvas Technologies (SHIELD, LifeCanvas Technologies). At 7 dpf, zebrafish were collected into Eppendorf tubes, 25 zebrafish per tube and anesthetized on ice briefly. Embryo buffer E3 was removed and replaced with 1 mL SHIELD Perfusion Solution with 4% paraformaldehyde (PFA) (Electron Microscopy Sciences), shaking overnight at 4°C. Whole zebrafish brains were dissected the next day and placed into tubes with fresh SHIELD Perfusion Solution, shaking overnight at 4°C. Tissue was placed into 1 mL SHIELD OFF solution, shaking overnight at 4°C. Tissue was transferred into SHIELD ON Buffer, shaking overnight at 37°C in MaxQ 4450 (ThermoFisher Scientific). Tissue was then cleared with 1 mL passive clearing protocol using SDS Clearing Solution, shaking for 5 days at 45°C. Clearing solution was washed off with 1 mL PBS + 1% Triton-X (PBT) with 0.02% Sodium Azide 3 times over 24 hours shaking at 37°C. Tissue was blocked in 1 mL PBT + 1% BSA for 2 hours shaking at room temperature, then incubated in primary antibody, shaking overnight at 4°C. Antibodies: 1:00 Synaptotagmin-1 (Lifespan Bioscience), GAD65 + GAD67 (Abcam), Beta-Actin (Abcam), 1:500 DAPI (Life Technologies), in 0.5 mL PBT + 1% BSA. Primary antibody was washed off 3 times in PBT and incubated in secondary antibody shaking overnight at 4 °C (1:500 488-555-680-Donkey Anti-Rabbit (Jackson ImmunoResearch) in PBT + 1% BSA). Secondary was washed off 3 times in PBT, then 1 mL EasyIndex was added to tissue, shaking overnight at room temperature. Whole brains were mounted in fresh EasyIndex on slides, placing coverslip with vacuum grease. Imaging was performed on an inverted Zeiss LSM700 Laser Scanning Confocal and processed on Fiji (ImageJ).

### Immunocytochemistry

Patient derived neurons were washed with PBS and fixed in fresh 4% paraformaldehyde in PBS overnight rocking at 4°C. Cells were washed with PBT and blocked in PBT + BSA for 1 hour at room temperature. Primary antibody was added to PBT overnight rocking at 4°C. Antibodies: 1:100 Synaptotagmin-1 (Syt1, Lifespan Bioscience), Vesicular Glutamate 1 and 2 (VGlut1/2, Synaptic Systems) or Postsynaptic Density 95 (PSD95, Abcam), Acetylated-Tubulin (Ac-Tubulin, Abcam). Primary antibody was washed off 3 times in PBT and incubated in secondary antibody shaking overnight at 4°C (1:500 488-555-680-Donkey Anti-Rabbit (Jackson ImmunoResearch) in PBT + BSA). Secondary was washed off 3 times in PBT, rocking for 2 hours at room temperature. Cells were washed 3 times in PBT and mounted on slides with DAPI (Prolong Gold Antifade with DAPI (Life Technologies). Imaging was performed on an inverted Zeiss LSM700 Laser Scanning Confocal and processed on Fiji (ImageJ).

SH-SY5Y cells were plated on coverslips and differentiated over 4 days with retinoic acid medium. The same imaging protocol was performed as above. Antibody: 1:200 Beta-Actin (Abcam).

### Western blot

HEK293 studies -Proteins were separated by SDS-PAGE and transferred to nitrocellulose membranes. HA-tagged constructs were identified using antibodies against HA or Flag peptides (1:5,000), and goat anti-rabbit or mouse horseradish peroxidase (1:10,000) were used as secondary antibodies (Jackson). Equal loading was confirmed using a mouse anti-GAPDH antibody. Detection was performed using the ECL detection system.

Larvae brain tissue (25 larvae brains pooled per sample) or differentiated SH-SY5Y cells (1 × 10^6^ cells per sample) were washed with PBS then lysed in RIPA buffer (Thermo) with protease inhibitor cocktail with a pestle homogenizer. Tissue/cells were rotate at 4°C for 30 min, then spun full speed 10 min. The supernatant was removed containing proteins, with denature in laemmli buffer for 1 hr at RT. Protein was separated on 10-40% gel and transferred PVDF by wet transfer. Membranes were blocked in 5% dry milk in TBS + Tween 20. Primary antibody was incubated overnight. Same antibodies were used for immunofluorescence and western analysis. Antibodies: Syt-1, Beta-Actin, GAPDH (Abcam). Secondary antibodies 1:2000 IRDye (Li-Cor) were incubated for 1 hour at room temperature. Blots were imaged and quantified on a Li-Cor Odyssey.

### Biotinylation and MS/MS

At 7 dpf, larvae were deeply anesthetized with tricaine. Larvae were dissected in PBS with protease inhibitor cocktail on ice, pooling 20 brains per genotype per sample. Assay was performed according to protocol utilizing Pierce Cell Surface Protein Isolation Kit (Thermo Fisher) with the following modifications. 1 vial of biotin was resuspended in 2 mL PBS and fresh brains were incubated with 1 mL biotin/PBS solution rotating for 45 minutes at 4C. Elution of biotin-bound proteins in water + DTT 1 hour at room temperature. Eluates were reduced, alkylated and digested with trypsin at 37°C overnight. This solution was subjected to solid phase extraction to concentrate the peptides and remove unwanted reagents. Solution was injected onto a Waters NanoAcquity HPLC equipped with a self-packed Aeris 3.6 µm C18 analytical column 0.075 mm by 20 cm, (Phenomenex). Peptides were eluted using standard reverse-phase gradients. The effluent from the column was analyzed using a Thermo Orbitrap Elite mass spectrometer (nanospray configuration) operated in a data dependent manner for 54 minutes. The resulting fragmentation spectra were correlated against the known database using Mascot (Matrix Science). Scaffold Q+S (Proteome Software) was used to provide consensus reports for the identified proteins. PEAKS Studio 8.5 was used for data analysis as a supplement to Mascot.

### Gene Ontology Analysis

GO SLIM analysis was performed with PANTHER Classification System (www.pantherdb.org) that combines genomes, gene function classifications, pathways and statistical analysis tools to enable biologists to analyze large-scale genome-wide experimental data^82^.

### Electrophysiology

iPSC Differentiated Neurons −1 × 10^4^ NPCs were plated and matured over 1 month in a 48-well CytoView plate (Axion Biosystems). Recordings of spontaneous activity were taken over 10-minute periods on the Maestro system (Axion Biosystems). AxIS software compiled the data collected from recordings. Data were collected for LFPs (firing frequency in Hz), electrographic burst events (minimum 5 LFPs/100 ms) and relative network activity (minimum 3 LFPs detected simultaneously between a minimum of two electrodes). LFP detection was filtered at 6 × standard deviation to remove potential artifacts. The external physiological solution contained (in mM) 128 NaCl, 5 KCl, 2 CaCl_2_, 1 MgCl_2_, 25 HEPES and 30 glucose, pH 7.3, Osmolarity 315 −325. The High KCl solution contained (in mM) 63 NaCl, 70 KCl, 2 CaCl_2_, 1 MgCl_2_, 25 HEPES and 30 glucose, pH 7.3, Osmolarity 315 −325.

Live larvae MEA recordings were performed as detailed in Tomasello and Sive (2020)^43^. For these recordings, larva was immersed in agarose in 12-well 64 electrode Cytoview plates (Axion Biosystems).

### Larval Behavior

At 7 dpf, dishes containing larvae were moved to the bench to allow acclimation to RT. With a cut 200 µl tip, larvae were individually pipetted into 96-well plates with 200 µl E3 media and moved to the Noldus Daniovision for 10 min habituation period. The larvae were exposed to a testing period of 70 minutes, with light (at 10%) extinguished for 5 seconds at 10-minute intervals. Point tracking collected distance and velocity traveled. Distance moved was calculated using the Ethovision XT 11 software from Noldus.

The same method is performed as above up to habituation. Baseline activity was then recorded for 10 minutes, followed by exchange of 100 µl E3 from each well with 100 µl of varying concentrations of PTZ to test a range of doses. Plates were immediately placed back on the Daniovision system for another 10 min recording. Point tracking collected distance and velocity traveled. Distance moved was calculated using the Ethovision XT 11 software from Noldus, normalizing to habituation time.

## Supplementary Methods

### PH3 Staining

At 24 hpf, *fam57b mut* and AB embryos were dechorionated and fixed overnight at 4°C in paraformaldehyde. Embryos were washed with phosphate buffered saline with Tween-20 (PBT) and yolk sac was removed. Embryos were incubated with 10% H_2_O_2_ for 1.5 hrs, then washed in PBT. Embryos were blocked in PBT with bovine serum albumin at room temp for 4 hrs, then incubated with α-PH3 antibody (1:1000, Upstate Biotechnology) overnight at room temp. Embryos were washed with PBT and incubated with secondary antibody (1:500 goat α-rabbit IgG HRP, Abcam) in PBT overnight at room temp. Embryos were washed in PBT and flat mounted on glass slide with propidium iodide in glycerol. Imaging was performed on a confocal microscope.

### TUNEL Staining

Embryos were collected, fixed and processed as PH3 staining. Embryos were then dehydrated then rehydrated interchanging ethanol and PBT. Proteinase K (In vitrogen) was incubated in PBT on neutator, then rinsed in PBT. TdT labeling was followed per manufacturer’s instructions, ApopTag kit (Chemicon). α-DIG-AP (1:100, Gibco) was used to detect the DIG labeled ends. Embryos were washed in PBT and flat mounted on glass slide with propidium iodide in glycerol. Imaging was performed on a confocal microscope.

### Neuromuscular Junction Staining

Staining performed as described in McCammon *et al*. (2017)^16^.

### Larvae Head Measurements

Measurements were taken as described in McCammon *et al*. (2017)^16^.

## Acknowledgements

We thank Olivier Paugois for excellent fish care and lab management. Special thank you for the Whitehead Institute core facilities. Specifically, we would like to thank Wendy Salmon (Keck Imaging Facility) for help with confocal imaging, Eric Spooner (MS Core) for proteomic work, Dr. Caroline Lewis (Metabolomics Core) for lipidomics analysis, and Dr. George Bell (Bioinformatics and Research Computing) for analysis and help in compiling MS/MS data. We thank Dr. Mike Gallager for help analyzing the published FAM57B mutation from Satterstrom *et al*. 2019^71^. Dr. Monika Saxena for constructive and helpful edits to the manuscript. Masters student Adrianna Vandeuren for help maintaining the fish lines and Undergraduate Research Opportunity Position Allysa Allen for help in expanding iPSCs. Finally, we would like to thank you to colleagues in the Simons Center for the Social Brain at MIT.

## Author Contribution

DLT and HS designed the study, interpreted results and wrote the manuscript. AHF helped conceive, supervise, interpret and write the CerS study (**Fig. 3**). JK and YK performed the experiments in **Fig. 3**. JMM created the *fam57b mutant* zebrafish line. MM and RJ instructed on stem cell culture work and provided unaffected control iPSC (originally from Coriell Institute Biobank). DLT performed all other studies.

## Data Availability

The authors declare that all data supporting the findings of this study are available within the article and its supplementary information. Raw data, including proteomics and lipidomics results, are available on appropriate request from corresponding author.

